# Monounsaturated fat prevent, whereas saturated fat promote, cholesterol crystals in cytoplasmic lipid droplets

**DOI:** 10.64898/2026.11.02.763728

**Authors:** Uche Njoku, Sierra Siedlecki, Lei Li, Amanda Quirk, Scott B Widenmaier

**Affiliations:** Department of Anatomy, Physiology, and Pharmacology, University of Saskatchewan, Saskatoon, Saskatchewan, Canada; Canadian Light Source, University of Saskatchewan, Saskatoon, Saskatchewan, Canada

**Keywords:** cholesterol metabolism, monounsaturated fatty acid (MUFA), cholesterol crystal, lipid droplet, liver

## Abstract

**Objective:** Metabolic dysfunction associated steatohepatitis (MASH) may, in part, result from excessive storage of cholesterol in hepatic lipid droplets that transition into a crystalline state. The mechanism by which crystals form in lipid droplets of hepatocytes and other cells such as macrophage is incompletely understood. Recently, we showed cholesterol must be esterified for crystals to form in lipid droplets. As the physicochemical properties of saturated cholesteryl ester render it more susceptible to crystallize than unsaturated cholesteryl ester, we investigated whether lipid droplet crystals are composed of saturated cholesteryl ester.

**Method:** Lipid droplet crystals were measured in cholesterol-loaded Hep3B hepatocyte-like and RAW 264.7 macrophage-like cells co-treated with vehicle, monounsaturated fat, or agents that increase cellular saturated fat level. Lipidomics was used to quantify cholesteryl ester species. We also used Fourier Transform Infrared (FTIR) spectromicroscopy on livers of mice with MASH to determine the molecular identity of crystals in hepatic lipid droplets.

**Results:** Cholesterol loading increased cholesteryl ester content and lipid droplet crystals. Monounsaturated fat treatment (e.g., oleate) increased the ratio of unsaturated to saturated cholesteryl ester and this corresponded with a substantial reduction in lipid droplet crystals. Treatments increasing saturated fat level did not exhibit this effect. (FTIR) spectromicroscopy identified cholesteryl palmitate as the prominent molecular constituent of hepatic lipid droplet crystals in mice with diet-induced MASH.

**Conclusion:** Hepatic lipid droplet crystals in mice with diet-induced MASH are composed of saturated cholesteryl ester and unsaturated fat intake supports the cellular capacity to sustain non-crystalline liquid disordered cholesteryl ester in lipid droplets.

## 1. Introduction

Metabolic dysfunction associated steatohepatitis (MASH) is a major health concern that increases the risk for cirrhosis, hepatocellular carcinoma, and cardiovascular disease [1, 2]. MASH occurs when simple steatosis progresses to a persistent state of liver injury, inflammation, and fibrosis [3, 4]. While there is incomplete understanding of its pathogenesis, mounting evidence suggest that the transition from simple steatosis to MASH is driven by the accumulation of lipids that exert toxic effects on hepatocyte viability and alter its structure and composition [4–7]. One lipid of growing interest is cholesterol [3].

Although cholesterol is required for homeostasis, excess cholesterol in the liver distinguish MASH from simple steatosis in humans [8–12] and rodent disease models [13–20]. This may be due to multiple mechanisms [3, 21, 22]. Elevated cholesterol in hepatic lipid droplets has been shown to play a major role and this may underlie the elevated risk for MASH in people with the *PNPLA3* I148M gene variant [23]. The mechanism linking excess cholesterol in lipid droplets to MASH is unclear but emerging evidence suggest it is triggered when cholesterol undergoes a physicochemical transition to a crystalline state [3]. Ioannou and colleagues showed that excess cholesterol promote cholesterol crystals in hepatic lipid droplets and that these crystals associate with MASH, but not simple steatosis, in humans and in mice [11, 12, 15, 16]. These studies reveal that cholesterol crystal-induced changes in hepatic lipid droplet composition, structure, or function may underlie MASH and its progression to severe liver disease states [3, 4, 6, 24].

Determining the mechanism underlying hepatic lipid droplet cholesterol crystal formation is critical to understand the role such crystals can play in MASH. Fundamentally, cholesterol crystals reflect the biophysical problem that cholesterol is prone to form an ordered crystal structure when its concentration exceeds the solubilizing capacity of its environment (e.g., cell membrane). Most cells have adaptive mechanisms to guard against this stressor [13, 20, 21, 25–27]. One is the capacity to produce cholesteryl ester using cholesterol and fatty acid substrates via sterol O-acyltransferase (SOAT), also known as acyl-coenzyme A: cholesterol acyltransferase [28]. The cholesteryl ester is then stored alongside triglyceride in lipid droplets. While often recognized as a safe storage site for excess cholesterol [26, 28], recent studies suggest this is not always the case. In cultured cells, increasing the proportion of cholesteryl ester to triglyceride has been found to alter lipid droplet organization and trigger a liquid crystalline state [29–32]. Moreover, using Raman scattering microscopy on lipid droplets in liver from humans with MASH, Xu et al. [12] spatially mapped triglyceride, cholesterol, and cholesteryl ester with a saturated or unsaturated fatty acid, and overlayed this with birefringence images that detect crystals. They found a higher spatial correlation between the crystals and heterogeneously organized saturated cholesteryl ester [12] than the correlation with cholesterol, unsaturated cholesteryl ester, and triglyceride, which is consistent with work on non-hepatic cells [29]. This suggests that hepatic lipid droplet crystals in humans with MASH may be composed of saturated cholesteryl ester. Moreover, using cell culture and mouse models we have shown that SOAT-dependent cholesterol esterification is required to form crystals in lipid droplets [33]. These studies indicate there are context-dependent limitations with respect to the storage capacity of lipid droplets for cholesteryl ester that contribute to the formation of MASH-associated cholesterol crystals.

Based on above, we hypothesize that saturated cholesteryl esters drive the formation of cholesterol crystals in cytoplasmic lipid droplet. Interestingly, it has been shown that monounsaturated fats (MUFA) such as oleate promote lipid droplet expansion and protect against cytotoxicity, whereas saturated fats such as palmitate promote cytotoxicity and facilitate formation of ordered structures [29–31, 34–38]. As the phase behavior of cholesteryl ester can depend on fatty acyl chain saturation, with saturated species more prone to crystallize at physiological conditions [38], we anticipate that increasing levels of unsaturated cholesteryl ester may prevent formation of lipid droplet crystals. Using cholesterol loaded Hep3B and RAW 264.7 cell models, here we investigate whether exogenous delivery of MUFAs protect against cholesterol crystals. Moreover, we utilize Fourier Transform Infrared spectromicroscopy on livers from mice with MASH to biochemically fingerprint cholesterol crystals within hepatic lipid droplets. Overall, our study shows that hepatic lipid droplet crystals in mice with diet-induced MASH are composed of saturated cholesteryl ester and suggests that the cellular capacity to sustain non-crystalline liquid disordered cholesteryl ester in lipid droplets is influenced by the ratio of saturated to unsaturated fatty acid. Consequently, our study indicates that boosting this adaptive capacity may mitigate the formation of MASH-associated cholesterol crystals.

## 2. Materials and methods

### 2.1. Cell culture and reagents

Human hepatoma Hep3B cells (ATCC, catalog #HB-8064) and RAW 264.7 cells (catalog #TIB-71) were purchased from American Type Culture Collection and maintained under standard culture conditions using a humidified incubator that was maintained at 37 °C and CO₂ level of 5 %, as we previously describe [33]. Treated cells were cultured on a 3.5 cm glass-bottom poly-D-lysine coated dish from Mattek (catalog #P35GC-1.5-14-C) in Dulbecco’s Modified Eagle Medium (DMEM) from Gibco (catalog #11965-092) and supplemented with 2.5 % cosmic calf serum from Cytiva Hyclone (catalog #SH30087.03).

Sources of reagents and consumables are as follows. Cholesterol (catalog #3045), cholesteryl oleate (catalog #C9253), cholesteryl palmitate (catalog #C6072), glyceryl trioleate (catalog #T7140), methyl-β-cyclodextrin (catalog #C4555), oleate (catalog #O1383), 16:0-18:1 phosphatidylethanolamine (catalog #850757P), 16:0-18:1 phosphatidylcholine (catalog #850457C), and sodium palmitate (catalog #P9767) were from Millipore Sigma. Palmitoleate (catalog #HY-W011873) and stearoyl-CoA desaturase-1 (SCD1) inhibitor A939572 (catalog #HY-50709) were from MedChemExpress. Dimethyl sulfoxide (DMSO; catalog #BP231-1), which served as vehicle control, and optical-bottom, black-wall 96-well plates (catalog #1256670) were from Fisher. Acetylated LDL (acLDL) was from Kalen Biomedical (catalog #770201-4). Hoechst 33342 (catalog #62249), BODIPY 493/503 (catalog #D3922), Amplex Red Cholesterol Assay Kit (catalog #A12216), and Pierce BCA assay kit (catalog #23225) were from ThermoFisher.

### 2.2. Preparation of the cholesterol-methyl β cyclodextrin (-MβCD) complex

As we described previously [33], cholesterol stock was generated by dissolving cholesterol powder (386.65 g/mol; density 1.067 g/mL at 25 °C) in 100 % ethanol at 70 °C to a final concentration of 100 mM. 1 ml aliquots of this stock were gradually added to 42 mg/mL MβCD (1320 g/mol) in deionized water with constant heating at 70 °C until reaching a final concentration of 5 mM cholesterol. Then, the cholesterol-MβCD complexed solution was slowly cooled to room temperature and left overnight at 4 °C to stabilize. The following day, it was sterile filtered and stored at 4 °C until use. This complex is referred to as “cholesterol” and has been previously validated as an efficient method for delivering excess levels of cholesterol into cultured cells [20, 33].

### 2.3. Imaging, detection, and quantification of cholesterol crystals and lipid droplets in cultured cells

4 × 10⁵ Hep3B cells or 1 × 10⁵ RAW 264.7 cells were seeded in 3.5 cm glass-bottom poly-d-lysine coated dishes and allowed to adhere overnight. The formation of cholesterol crystals was induced in Hep3B cells upon treatment with 50 μM or 200 μM cholesterol, whereas crystals were induced in RAW 264.7 cells upon treatment with 50 μg/mL acLDL. As indicated in the text and figures, the cells were co-treated with 50-400 μM of MUFAs, palmitoleate or oleate, or the saturated fatty acid, palmitate and an inhibitor of the fatty acid desaturation enzyme, stearoyl-CoA desaturase 1 (SCD1). Treatment containing media was refreshed every 24 hours. At the endpoint, media was removed and cells incubated in phosphate buffered saline (PBS) containing 5 μg/ml Hoechst 33342 for 15 minutes at 37 °C to stain nuclei. In experiments in which lipid droplets were analyzed, cells were also incubated with 2 μM BODIPY 493/503 to stain lipid droplets. Afterward, the media was replaced with fresh PBS.

Cholesterol crystal images in live cells were visualized via polarized light microscopy on an Olympus DP73 camera at 20 × magnification. Parallel fluorescence images of Hoechst 33342 stained nuclei (blue filter cube – ET DAPI) were collected to normalize crystal area to the number of imaged cells. As indicated, images of BOPIDY 493/503 stained lipid droplets (green filter cube – FITC-A-Basic-OFF) were collected to assess the relationship between lipid droplets and cholesterol crystals. Images were acquired from three non-overlapping fields per dish, with field selection initially performed in bright-field mode prior to crystal visualization to avoid sampling bias. All images within a given experiment were obtained under uniform exposure conditions across channels.

Abundance of crystals and lipid droplets were quantified in cultured cells using ImageJ software (https://imagej.nih.gov/ij/) as previously described [33], with slight modifications. For crystals, raw images were converted from RGB to 8-bit format and the pixel threshold was set between 20-255 for Hep3B cells and between 15-255 for RAW 264.7 cells. Then we measured the % crystal area. For lipid droplets, raw images were converted to 8-bit format and the pixel threshold was set between 20-255 to isolate lipid-positive structures. The background was set to black and threshold image was made into a binary mask. A watershed was applied to isolate and measure individual lipid droplets. Particles were analyzed with the size range set 0-infinity and circularity set between 0.00-1.00. Then we measured the image and recorded the number and area of lipid droplets.

To determine cell number per field of view, corresponding Hoechst 33342 stained nuclei images were converted from RGB to 16-bit format. The pixel threshold was set between 10-65535 for Hep3B nuclei and between 15-65535 for RAW 264.7 nuclei. Next, images were processed using the watershed function to separate overlapping nuclei into single objects. Particles were analyzed with the size range set 3000-infinity for Hep3B cells and 2000-infinity for RAW 264.7 cells, with circularity set between 0.00-

1.00. Objects were quantified to determine cell number, and crystal or lipid droplet abundance was normalized to cell number and expressed as % positive area or number per 100 cells, as indicated in figures and text. The mean of three crystal or lipid droplet images per dish was considered to represent one biological replicate and is shown in results as a single data point, with each biological replicate representing an independent experiment.

### 2.4. Quantification of cholesterol crystals in mouse liver sections

Hepatic cholesterol crystals in mouse liver sections were analyzed as previously described [33]. Briefly, optimal cutting temperature compound (OCT) embedded liver tissue was snap frozen on dry ice, cryo-sectioned at 10 mm thickness and mounted in glycerol. Cholesterol crystals were visualized via polarized light microscopy and quantified using ImageJ as done in section 2.3, except in this case the crystal abundance was expressed as % of crystal positive area in total image of liver section.

### 2.5. Lipidomic Analyses of cultured Hep3B cells

8 × 10^6^ Hep3B cells were seeded in 150 mm culture plates. Cells were allowed to grow to 60 % confluency before undertaking 48 hours of treatment. At the endpoint, media was removed and cells were washed twice with ice-cold PBS. Then, cells were scraped and collected from the plate, resuspended in 1 ml ice-cold PBS, and underwent centrifugation at 500 g and 4**°**C for 10 minutes. The supernatant was removed, and the cell pellet was allowed to dry on ice for 5 minutes before being weighed and snap frozen by a liquid nitrogen plunge. Samples were stored at −80 **°**C until being sent to The Metabolomics Innovation Center (https://metabolomicscentre.ca) at the University of Alberta for Ultra High-Performance Liquid Chromatography coupled to High Resolution Mass Spectrometry (UHPLC-HRMS), utilizing the global lipidomic platform as well as the bioinformatic and statistical analysis.

### 2.6. *In vivo* mouse study and liver collection

*In vivo* procedures were approved by the University of Saskatchewan’s Animal Care Committee. Male and female C57bl/6J mice from the Jackson Laboratory (www.jax.org/strain/000664) were housed at 21 °C on a 12-hour light/dark cycle and provided *ad libitum* access to food and water. Beginning at 8-10 weeks of age, mice were fed for 24 weeks with regular chow (Prolab RMH 3000 from LabDiet) or with a MASH-inducing high fat, fructose, and cholesterol (HFFC) diet (Research diets, catalog# D09100310) enriched with 40 % fat (75 % of which is saturated fat rich palm oil), 20 % fructose, and 2 % cholesterol. As we have shown previously [13, 33, 39], mice fed HFFC diet for >16 weeks develop features of MASH, including hepatic lipid droplet cholesterol crystals. At the endpoint, mice were euthanized under anesthesia with 3 % isoflurane at an oxygen flow rate of 1 L/min. The vasculature was flushed with 20 ml of room temperature PBS. A portion of liver was snap frozen and stored at −80 °C. A second portion from the left lobe was fixed in 10 % neutral buffered formalin, embedded in wax, sectioned at 5 μm thickness, and stained with Hematoxylin and Eosin (H&E). A third portion was embedded in OCT compound, frozen on dry ice, and cryo-sectioned at a 10 μm thickness, and this was used for assessing cholesterol crystals in liver or for identifying the biochemical fingerprint within these crystals (described further in section 2.8).

### 2.7. Fourier Transformed Infrared (FT-IR) Spectromicroscopy and Data Processing

Liver sections were placed on calcium fluoride (CaF_2_) windows (Crystran Ltd, UK) and FTIR spectromicroscopic images were collected at the Mid-infrared beamline (01B1-01) at the Canadian Light Source (Saskatoon, Saskatchewan, Canada, https://www.lightsource.ca/index.php). Hyperspectral images were collected with the thermal glowbar source of the Agilent endstation, which consists of an Agilent Cary 760 spectrometer with a Cary 620 Microscope and a liquid nitrogen-cooled 128 pixel × 128-pixel Focal Plane Array (FPA) detector. The measurements were made using a 25 × magnification objective and matching condenser lenses with a numerical aperture of 0.81. Spectra were collected in transmission mode over the spectral range of 3900–900 cm⁻¹ at a spectral resolution of 4 cm⁻¹ using 32 co-added scans and 64 co-added background scans and a pixel size of 3.3 μm. A background spectrum was collected from a clean CaF_2_ window before sample acquisition. Spectral preprocessing and analysis were done using Quasar software (1.1.1 [40]). All spectra were corrected for atmospheric water vapor and CO₂ contributions, and baseline corrected. Spectral maps were generated from integrated peak intensities unless otherwise stated.

### 2.8. Statistical Analysis

Data were analyzed using GraphPad Prism software (version 9.5.1). Statistical significance was set at *p* < 0.05. Depending on the experimental design, comparisons were evaluated using unpaired *t*-tests, simple linear regression, or one-way and two-way analysis of variance (ANOVA) followed by Dunnett’s or Tukey’s post hoc tests, as specified in the corresponding figure legends. Results are presented as mean ± standard error of the mean (SEM). All experiments were performed with at least three independent biological replicates, and the number of samples included in each analysis is reported in the figure panels.

## 3. Results

### 3.1. Excess MUFA protects against cholesterol crystals in cholesterol-loaded Hep3B cells

We previously showed that treating the hepatocyte cell line, Hep3B cells, with 200 μM cholesterol for 48 h results in the formation of cholesterol crystals in lipid droplets and that this process requires the esterification of cholesterol [33]. Continuing with this experimental model, we investigated whether cholesterol-induced crystal formation is affected by excess monounsaturated fatty acid (MUFA) exposure, which Hep3B cells can use to produce cholesteryl ester or triglyceride and then store in lipid droplets.

First, we tested the effect of treating with the 18 carbon MUFA, oleate. The workflow is shown in Fig. 1A. Hep3B cells were treated with 200 μM cholesterol in the presence or absence of 200 μM or 400 μM oleate, with bovine serum albumin (BSA) used as control. As expected, oleate treatment increased triglyceride content in lipid droplets, as detected via fluorescence imaging of BODIPY (Fig. 1B). Polarizing light microscopy was used to detect birefringent cholesterol crystals, as previously done [33]. Fluorescence detection of Hoechst 33342 stained nuclei was used to normalize the abundance of crystals or lipid droplets by the number of cells per image. As shown in Fig. 1B and C, crystal formation did not occur in cells treated with only BSA or 400 μM oleate whereas cells treated with 200 μM cholesterol had a robust level of crystals and relatively similar lipid droplet content compared to control cells. In contrast, cholesterol crystals were significantly reduced in cholesterol-loaded Hep3B cells when they were co-treated with 200 μM or 400 μM oleate, corresponding with a substantial increase in lipid droplet level (Fig. 1B and C). To determine if this effect was specific to oleate or a general effect of MUFAs, we replicated this experiment using the 16 carbon MUFA palmitoleate. Like oleate, palmitoleate substantially reduced cholesterol crystal formation and increased lipid droplets in the cholesterol loaded Hep3B cells (Fig. S1A and B). This evidence shows that excess MUFA prevents cholesterol-induced lipid droplet crystal formation in Hep3B cells.

**Figure 1.**
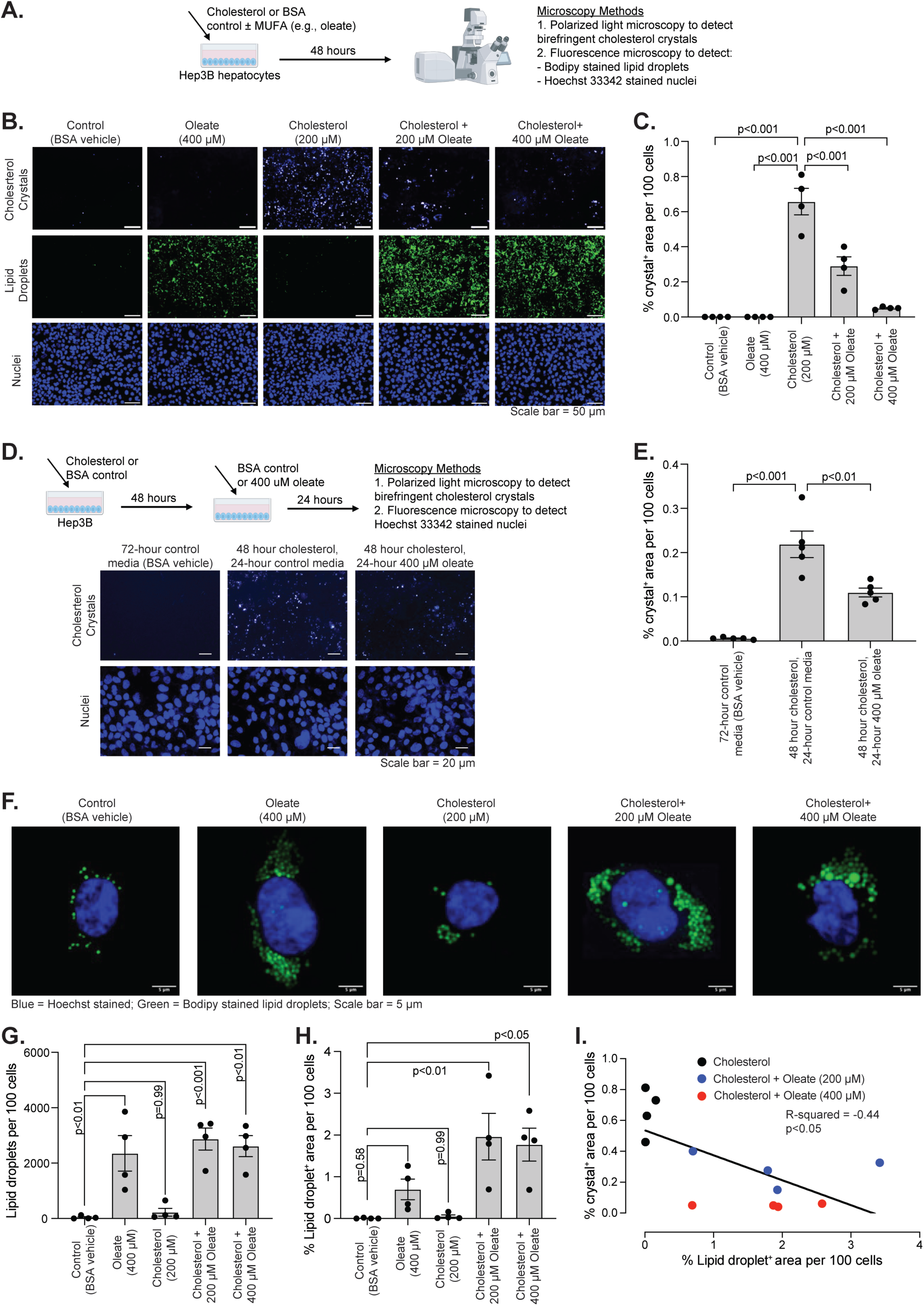
Supplementation with the monounsaturated fat oleate prevents formation of cholesterol crystals in Hep3B cells. (A) Study design. (B) Representative images of cells treated without or with 200 μM cholesterol ± 200 μM or 400 μM oleate. Identified crystals, nuclei, and lipid droplets are shown in panel. (C) Quantified crystals in (B) to examine the effect of oleate on cholesterol crystal formation (n = 4). (D) Study design and representative images of cells treated without or with cholesterol for 48 h and then media replaced with indicated media for 24 h. (E) Quantified crystals in (D) to examine the effect of oleate in resolving cholesterol crystals (n = 5). (F) Representative higher magnification images of cells treated as indicated in panel. (G-H) Quantified number (G) and area (H) of lipid droplets (n = 4). Data in C, E, G, and H are mean ± standard error of the mean and the *p*-values were determined by one-way analysis of variance, with Tukey’s post-hoc test. (I) Correlation analysis of cholesterol crystals with lipid droplets area. The individual data points and slope line are shown (n = 12). The *p*-value and slope were determined using simple linear regression analysis.

Previously [14], we showed crystals that form in Hep3B cells following 48 h of 200 μM cholesterol treatment will dissipate by ≈ 50 % after switching the cells back to control media for another 48 h. In other words, Hep3B cells can resolve cholesterol crystals in control media. To investigate whether exogenous oleate accelerates crystal resolution, we undertook the experimental workflow shown in Fig. 1D. Control cells were treated with BSA for 48 h and then BSA containing media was replaced for another 24 h, resulting in no detectable crystals (Fig. 1D and E). In contrast, the other two groups were treated with 200 μM cholesterol for 48 h and then cholesterol containing media was replaced with either BSA containing media or media containing 400 μM oleate for another 24 h. Compared to BSA media, oleate significantly reduced the level of crystals in these cholesterol-loaded cells (Fig. 1D and E). This shows that, in addition to preventing crystals, MUFAs can also increase the resolution rate of pre-existing cholesterol crystals.

We next examined the relationship between lipid droplets and cholesterol crystals. The number and area of lipid droplets were more abundant in the cells treated with oleate (Fig. 1 F-H) and palmitoleate (Fig. S1C-E), likely reflecting increased triglyceride storage, and there was a significant inverse correlation with cholesterol crystals (Fig. 1I and S1F). Altogether, our results indicate that MUFA-loaded lipid droplets are protected against crystal formation in cholesterol-loaded Hep3B cells.

### 3.2. Excess MUFA protects against cholesterol crystals in cholesterol-loaded RAW 264.7 cells

To investigate whether MUFA also protects against lipid droplet cholesterol crystals in non-hepatic cells, we studied RAW 264.7 M1-like macrophages. Excess cholesterol uptake from modified low-density lipoprotein (LDL) has been shown to induce crystals in proinflammatory M1-like macrophages, an effect linked to atherosclerosis [26, 41]. Similar to Hep3B cells, we have shown [33] cholesterol crystals form in RAW 264.7 cells exposed to 50 μg/ml acetylated LDL for 48 h in a SOAT-dependent manner, indicating that the mechanism of lipid droplet crystal formation is similar despite being different cell types.

The workflow using RAW 264.7 cells in Fig. 2A is similar to the workflow on Hep3B cells, except in this case the cells were treated with acetylated LDL. As before [33], we found that acetylated LDL caused robust crystal formation whereas crystals were not detected in BSA control or oleate treated cells (Fig. 2B and C). Importantly, 50 μM or 400 μM oleate co-treatment reduced crystals in a dose dependent manner. Also, treatment with acetylated LDL + oleate increased the abundance and number of lipid droplets (Fig. 2B, D, and E), which correlated inversely with cholesterol crystals (Fig. 2F). These findings combined with the results in Figure 1 demonstrate that MUFAs can protect against cholesterol crystals in hepatocytes and macrophages, two cell types capable of storing large amounts of cholesterol.

**Figure 2.**
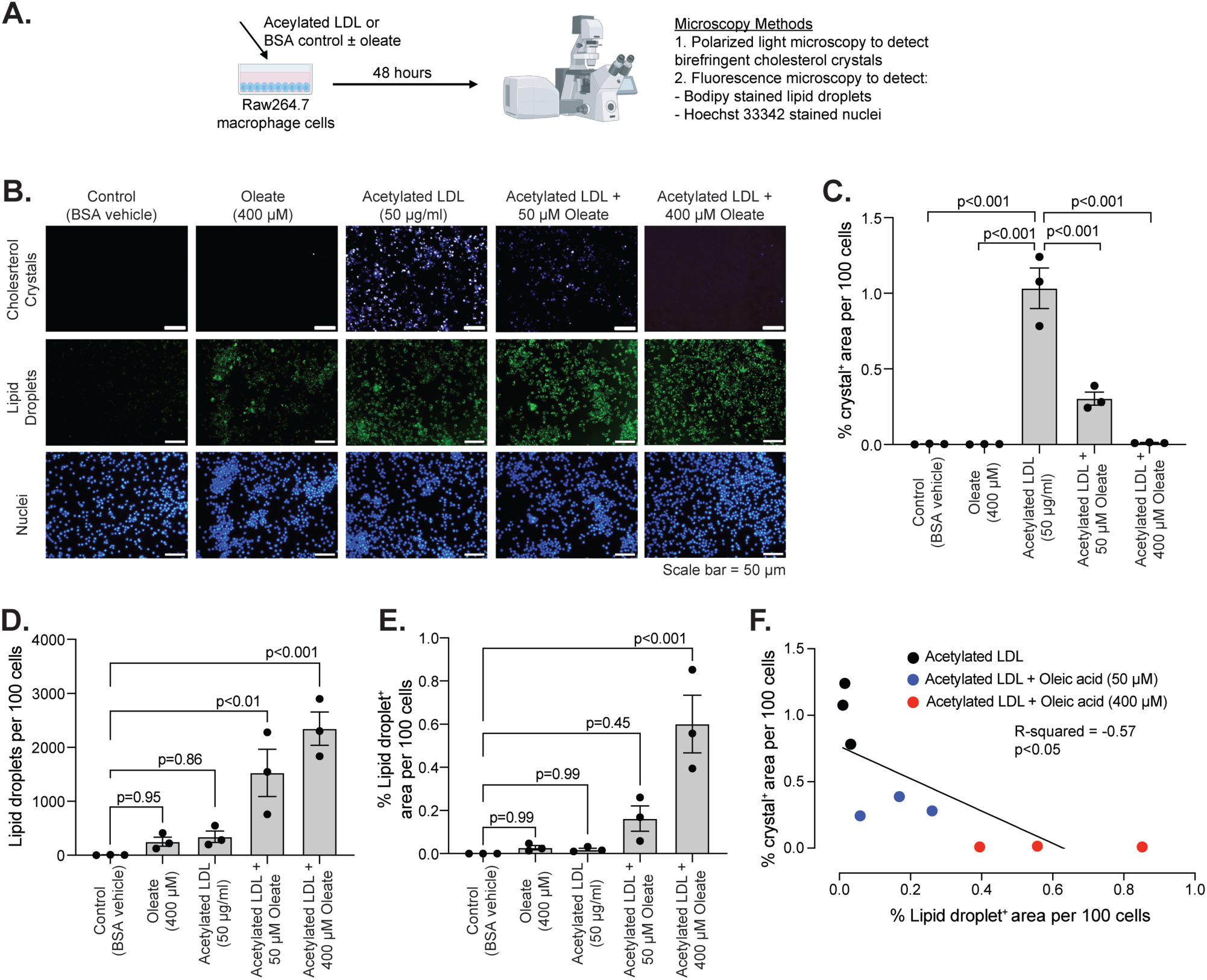
Supplementation with the monounsaturated fat oleate prevents formation of cholesterol crystals in RAW 264.7 cells. (A) Study design. (B) Representative images of cells treated without or with 50 μg/ml acetylated LDL ± 50 μM or 400 μM oleate. Identified crystals, nuclei, and lipid droplets are shown in panel. (C-E) Quantified crystals in (B) to examine the effect of oleate on cholesterol crystal formation as well as the number (D) and area (E) of lipid droplets (n = 3). Data in C-E are mean ± standard error of the mean and the *p*-values were determined by one-way analysis of variance, with Tukey’s post-hoc test. (F) Correlation analysis of cholesterol crystals with lipid droplets area. The individual data points and slope line are shown (n = 9). The *p*-value and slope were determined using simple linear regression analysis.

### 3.3. Effect of palmitate and inhibiting fatty acid desaturation on the formation of cholesterol crystals in Hep3B cells

Saturated cholesteryl ester can form ordered structures and have a greater tendency than unsaturated cholesteryl ester to transition into a crystalline phase [38]. While oleate treatment may prevent crystals by increasing unsaturated cholesteryl esters that are prone to remain in a liquid disordered state, treatment with the saturated fatty acid palmate may increase saturated cholesteryl esters to promote crystals. To investigate, we treated Hep3B cells with BSA, palmitate, cholesterol, or palmitate + cholesterol and examined the effect on the formation of cholesterol crystals. With palmitate however, using the dose and timing that match the oleate experiment (Fig. 1A-C) resulted in substantial loss of cultured cells. To maintain viable cells we adjusted the experimental condition to 200 μM palmitate for only 24 h. Though a shorter time frame, there was still a significant increase in crystals in cells treated with 200 μM cholesterol (Fig. 3A-B). Interestingly, there was a similar increase in crystals in the cells treated with cholesterol + palmitate, and palmitate on its own caused a trending rise in crystals. Our finding shows that in stark contrast to the effect of MUFAs, cellular exposure to excess palmitate does not prevent cholesterol crystals.

**Figure 3.**
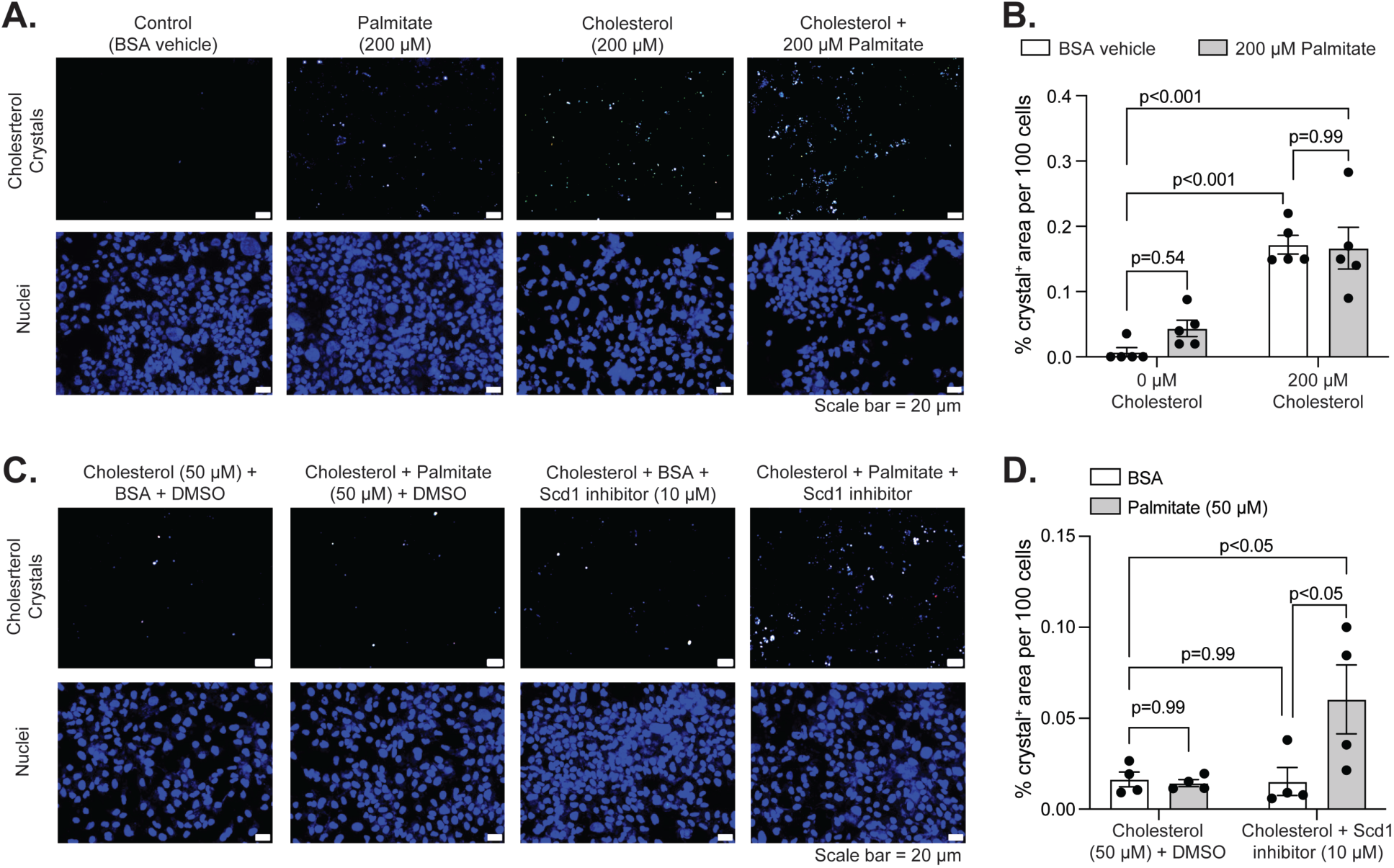
Treatments that increase cellular content of saturated fat does not prevent formation of cholesterol crystals in Hep3B cells. (A) Representative images of cells treated without or with 200 μM cholesterol ± 200 μM palmitate for 24 h. Identified crystals and nuclei are shown in panel. (B) Quantified crystals in (A) to examine the effect of palmitate on cholesterol crystal formation (n = 5). (C) Representative images of 50 μM cholesterol-treated cells also treated without or with 50 μM palmitate ± 10 μM SCD1 inhibitor for 24 h. Identified crystals and nuclei are shown in panel. (D) Quantified crystals in (C) to examine the effect of suppressing fatty acid desaturation on cholesterol crystal formation in cholesterol and palmitate-loaded cells (n = 4). Data in B and D are mean ± standard error of the mean and the *p*-values were determined by one-way analysis of variance, with Tukey’s post-hoc test.

Stearoyl-CoA desaturase 1 (SCD1) is an abundant hepatic enzyme that converts saturated fatty acid to MUFA and thus can limit saturated cholesteryl ester level in cells. We investigated if inhibiting SCD1 in cholesterol-loaded Hep3B cells exposed to a modest level of palmitate can increase cholesterol crystals. Hep3B cells were treated for 24 h with 50 μM cholesterol combined with DMSO, 50 μM palmitate, 10 μM SCD1 inhibitor (A939572), or 50 μM palmitate + 10 μM SCD1 inhibitor. We found the combination of cholesterol + palmitate + SCD1 inhibitor increased crystal levels greater than in cells receiving cholesterol only, whereas this effect did not occur in the cells receiving cholesterol + palmitate or cholesterol + SCD1 inhibitor (Fig. 3C and D). These findings show the saturated fatty acid palmitate has a markedly different effect on lipid droplet cholesterol crystal formation than the MUFAs, oleate and palmitoleate.

### 3.4. Effect of oleate on the lipidome and cholesteryl ester composition in Hep3B cells

To determine whether oleate may prevent cholesterol crystal formation by increasing the relative abundance of unsaturated cholesteryl ester, we examined the lipidome of Hep3B cells treated for 48 h with BSA control, 400 μM oleate, 200 μM cholesterol, or 200 μM cholesterol + 400 μM oleate. To do this, we used Ultra High-Performance Liquid Chromatography coupled to High Resolution Mass Spectrometry as well as a global lipidomic platform and related bioinformatic and statistical analysis available from The Metabolomics Innovation Center (https://metabolomicscentre.ca/). Principal Component Analysis (PCA) revealed reproducible within-group scores, with the profiles in cholesterol treated cells being more similar to control cells as compared to oleate treated cells (i.e., oleate or cholesterol + oleate), which exhibited a markedly distinct profile (Fig. 4A).

**Figure 4.**
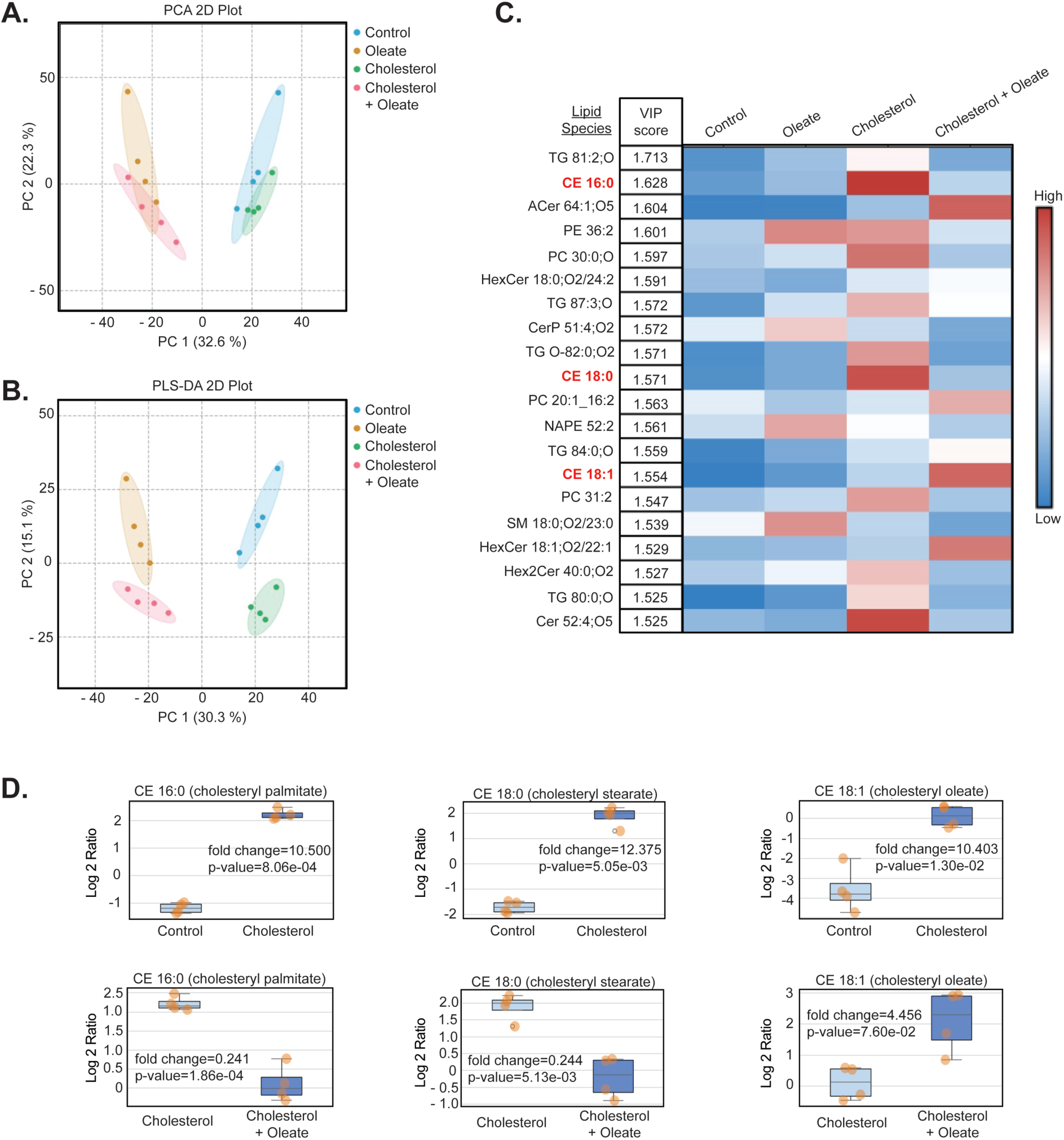
Lipidomic analysis show that oleate-induced lipid remodeling increase the ratio of unsaturated to saturated cholesteryl ester in cholesterol-loaded Hep3B cells. (A-B) A 2-dimensional Principal Component Analysis (PCA) in (A) and Partial Least Squares-Discriminant Analysis (PLS-DA) in (B) of lipid profiles for Hep3B cells treated with BSA, 400 μM oleate, 200 μM cholesterol, or 200 μM cholesterol + 400 μM oleate for 48 h (n = 4). (C) Heatmap showing relative change for the 20 top ranked lipids underlying group separation in (B), as determined by a PLS-DA Variable Importance in Prediction (VIP) score. Highlighted on the left in red are cholesteryl ester species (i.e., cholesteryl palmitate, cholesteryl stearate, and cholesteryl oleate) in this top 20 ranking. (D) Fold change of cholesteryl palmitate, cholesteryl stearate, and cholesteryl oleate comparing cholesterol versus control in the top graphs and cholesterol + oleate versus cholesterol in the bottom graphs (n = 4). The *p*-values were determined by univariate statistical analysis using NovaMT LipidScreener.

Partial Least Squares-Discriminant Analysis (PLS-DA) showed a similar outcome to the PCA plot (Fig. 4B). A PLS-DA Variable Importance in Prediction (VIP) score was generated to rank lipids according to the degree of impact on lipid profiles. The 20 most important are shown in Fig. 4C, labeled according to the LIPID MAPS nomenclature (https://www.lipidmaps.org/). Notably, the saturated cholesteryl esters cholesteryl palmitate (CE 16:0) and cholesteryl stearate (CE 18:0) as well as the unsaturated cholesteryl ester cholesteryl oleate (CE 18:1) were identified and the corresponding heatmap (Fig. 4C) and separate univariate analyses show that, as expected, each of these cholesteryl esters were increased in cells treated with cholesterol compared to the control cells (top graphs in Fig. 4D). In contrast, the levels of cholesteryl palmitate and cholesteryl stearate were reduced whereas cholesteryl oleate was increased in cells treated with cholesterol + oleate compared to cholesterol treated cells (bottom graphs in Fig. 4D). Our results show that oleate treatment caused the Hep3B cells to increase the relative proportion of unsaturated to saturated cholesteryl ester while storing excess cholesterol, an effect that may explain how excess oleate exposure prevents the formation of crystals in cholesterol loaded cells.

### 3.5. Identification of hepatic lipid droplet cholesterol crystals in liver of mice with HFFC diet-induced MASH using Fourier Transform mid-infrared (FTIR) spectromicroscopy

Our results using cholesterol-loaded Hep3B cells support the hypothesis that saturated cholesteryl ester can drive formation of hepatic lipid droplet crystals. Next, we turned our attention to a mouse model of MASH. Previously we demonstrated that mice fed for 24 weeks with a high fat, fructose, and cholesterol (HFFC) diet that contains enriched levels of palmitic acid and cholesterol will develop liver steatosis, inflammation, and fibrosis (i.e., MASH) as well as hepatic lipid droplet cholesterol crystals [13, 33, 39]. We used this *in vivo* model here to identify the dominant lipid component of hepatic lipid droplet crystals in the context of diet-induced MASH. Male and female C57BL/6J mice were fed regular (Control) diet or HFFC diet for 24 weeks, underwent euthanasia, and then livers were collected for examination (Fig. 5A). Compared to Control, HFFC diet fed mice had greater body (Fig. 5B) and liver (Fig. 5C) weight. Analysis of hematoxylin and eosin-stained livers sections (Fig. 5D) showed HFFC diet fed mice had substantially greater liver steatosis (Fig. 5E) and inflammation (Fig. 5F). Polarizing light microscopy on liver sections revealed a robust level of birefringent crystals in the HFFC diet fed mice (Fig. 5G) that correspond with a markedly greater crystal positive area compared to mice fed Control diet (Fig. 5H), which had virtually no detectable crystals. Hence, our findings show that HFFC diet fed mice developed cholesterol crystals and several other features resembling MASH [11, 12], like other models [15, 16].

**Figure 5.**
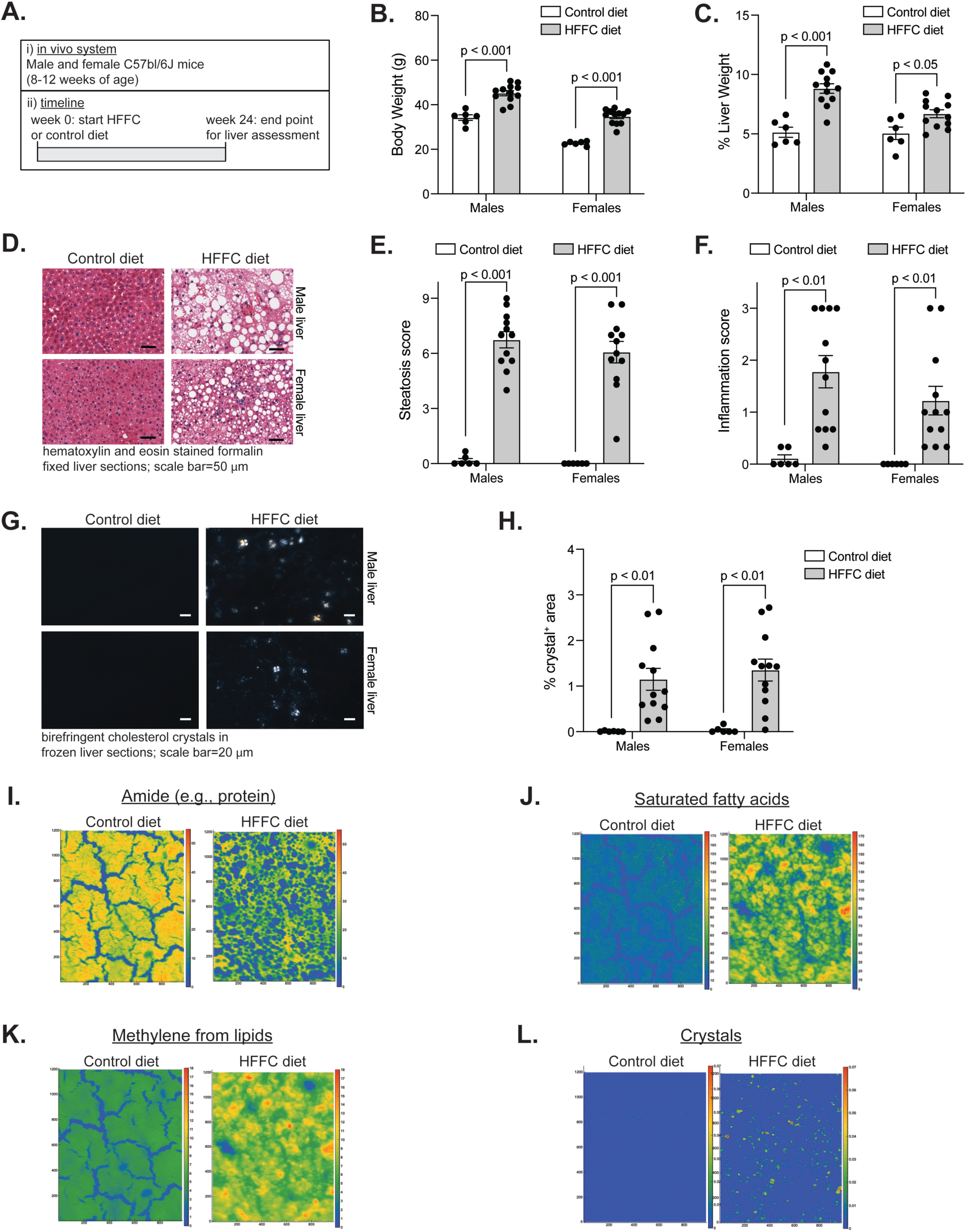
Detection of hepatic lipid droplet crystals in the livers of mice with HFFC diet-induced MASH using Fourier Transform Infrared (FTIR) spectromicroscopy. (A) Study design (n = 6 for control, n = 12 for HFFC). (B-C) Bodyweight (B) and % liver weight (C). (D-F) Representative images of hematoxylin and eosin-stained liver sections (D) and the steatosis (E) and inflammation (F) scores. (G-H) Representative images of crystals detected in the liver sections (G) and related quantification of % crystal area (H). (I-L) Representative spatial heatmaps of amide bonds (I), saturated fatty acids (J), methylene groups (K), and crystals (L). Data in B, C, E, F, and H are mean ± standard error of the mean, with individual data points shown. The *p*-values were determined by t-test, with adjustment for multiple comparison.

Next, we used Fourier Transform Infrared (FTIR) spectromicroscopy to compare the biochemical composition of livers from HFFC and Control diet fed mice. FTIR spectromicroscopy measure molecular vibrations within a molecule by assessing the absorption of infrared light at specific wavelengths [42]. The resulting spectrum provides a biochemical fingerprint of molecules within a tissue. In our samples, adjacent liver sections were used to detect birefringent crystals via polarized light microscopy and for full field FTIR imaging. The detailed biochemical maps enabled us to differentiate between crystal containing regions, lipid droplets, and surrounding tissue which was obtained from the high spatial resolution of FTIR imaging. In infrared spectroscopy, variations in the peak intensity, position, and spectral profile revealed key changes in composition and molecular organization. According to established methods [42–46], we analyzed spectra features corresponding to amide bonds (i.e., protein), total lipid, ester carbonyl groups (C=O), methylene (CH₂) and methyl (CH₃) stretching vibrations, olefinic (HC=CH) groups, and the cholesterol-associated vibrational modes. Heatmap images from integrated peak intensities show the relative difference in livers from Control versus HFFC diet fed mice with respect to the abundance of protein (Fig. 5I), saturated fatty acid (Fig. 5J), total lipid (Fig. 5K), and crystals (Fig. 5L). Critically, crystals were detected in all of the HFFC diet fed mice but not in any of the Control diet fed mice.

### 3.6. FTIR spectromicroscopy identifies cholesteryl palmitate as the molecular constituent of hepatic lipid droplet crystals in the livers of mice with HFFC diet-induced MASH

Once establishing the detection of hepatic lipid droplet crystals in the mice with HFFC diet-induced MASH, we used FTIR spectromicroscopy to determine the molecular identity of these crystals. Shown in Fig. 6A are RGB biochemical maps for relative distributions of protein-rich (green), lipid-rich (red), and crystal-containing (blue) regions in livers from Control and HFFC diet fed mice. The white arrows show selected areas of crystal containing regions, which are all within the lipid-rich droplets. The spectra of the crystal-containing regions was extracted from 25 crystals per liver section collected from 5 males and 5 females and compared with surrounding lipid droplets and reference lipid standards to identify the species. In crystal containing regions, the spectra exhibited an ester carbonyl peak at ≈1740 cm⁻¹ and distinct sharp CH_2_ wagging mode peaks between 1350 and 1180 cm⁻¹, indicative of ordered crystalline lipid packing [47]. These features were absent in the surrounding non-crystalline regions. To ascertain its molecular identity, the spectra profile of these crystals were compared to those of reference lipid standards. In Fig. 6B is unique spectra features for oleate, trioleate, cholesteryl oleate, cholesteryl palmitate, cholesterol, palmitate, and 1-palmitoyl-2-oleoyl-sn-glycero-3-phosphocholine that highlight the ester carbonyl peak (≈1740 cm⁻¹) and the distinct sharp wagging mode peak regions between 1350 and 1180 cm⁻¹. Notably, the spectra profile of cholesterol was clearly distinct from cholesteryl esters (Fig. 6B) whereas the wagging mode peaks between 1350 and 1180 cm⁻¹ were the key features distinguishing cholesteryl oleate and cholesteryl palmitate (Fig. 6C). Importantly, in the infrared spectra the CH_2_ wagging modes only arise from ordered long chain alkane molecules and we found that the spectra profile for the hepatic lipid droplet crystals (i.e., blue region in Fig. 6A) exhibited a sharp peak around 1740 cm⁻¹ as well as the CH_2_ wagging mode peaks between 1350 and 1180 cm⁻¹, which were at identical peak positions as compared to cholesteryl palmitate (left graph in Fig. 6D) and highly distinct from the peak positions for cholesteryl oleate (right graph in Fig. 6D). Hence, our findings show that hepatic lipid droplet crystals in mice with HFFC diet-induced MASH are composed of cholesteryl palmitate, consistent with recent findings in the liver from humans with MASH [12].

**Figure 6.**
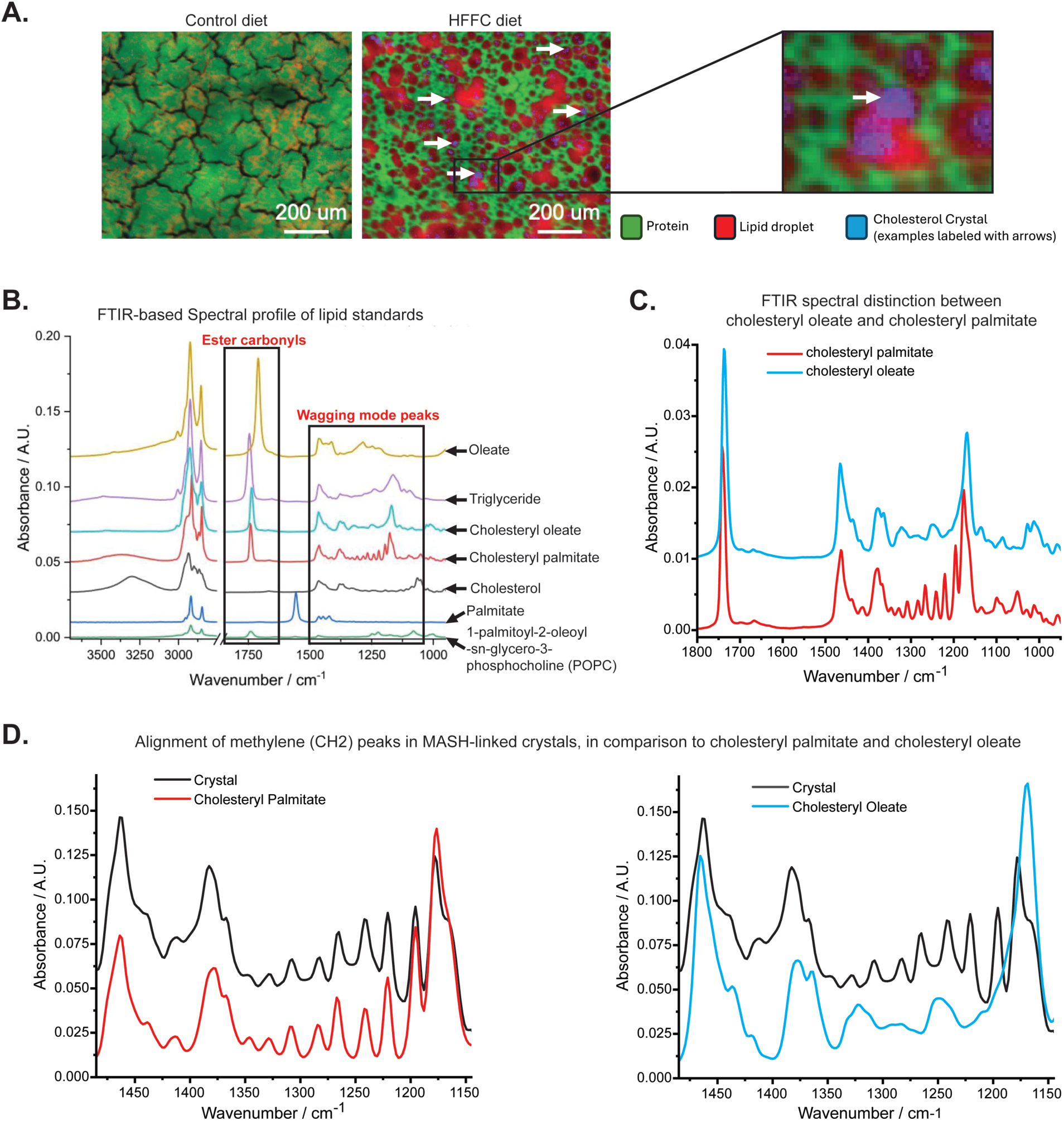
FTIR spectromicroscopy reveal that hepatic lipid droplet crystals in the livers of mice with HFFC diet-induced MASH is composed of cholesteryl palmitate. (A) RGB biochemical map showing the relative distributions of protein-rich (green), lipid-rich (red), and crystal-containing (blue) regions in the livers of Control and HFFC diet fed mice. White arrows show selected areas of crystal containing regions. Spectra was extracted from 25 crystals per liver to generate a spectra profile (n = 5 males and 5 females). (B-C) Spectra profile showing reference lipid standards and highlighting ester carbonyl and wagging mode peak regions (B) and comparing the spectra profiles (C) of cholesteryl palmitate versus cholesteryl oleate. (D) Comparisons of spectra profiles showing that hepatic lipid droplet crystals in the liver of mice with MASH have CH_2_ wagging mode peaks at identical peak positions as cholesteryl palmitate (left graph) and highly distinct from cholesteryl oleate (right graph).

## 4. Discussion

Metabolic dysfunction-associated steatohepatitis (MASH) is a progressive state of liver disease that promotes severe clinical pathologies and existing therapies are insufficient to address this global challenge [1–6]. Over the past two decades elevated hepatic cholesterol has emerged as a key factor underlying MASH [3, 8–10, 22]. While the mechanism for this is not understood, several studies suggest cholesterol in lipid droplets play a role. Sakuma et al. [23] showed that elevated hepatic lipid droplet cholesterol may mediate the onset of MASH and Ioannou and colleagues have established that hepatic lipid droplet cholesterol crystals are a distinguishing feature of MASH [3, 11, 15, 16, 18]. However, a well-known crystal sensing inflammasome mediator was found to be dispensable for crystal-associated MASH to develop in a dietary mouse model of the disease [48] whereas Li et al. showed that hepatic lipid droplet crystals directly stiffen the liver prior to fibrosis in a rat model of steatotic liver disease [49]. Consequently, it remains unclear if and how hepatic lipid droplet cholesterol crystals contribute to MASH and its progression.

Determining the mechanism by which crystals form in cytoplasmic lipid droplets may reveal the role they play in MASH and how to alleviate them. Previously, we showed that cholesterol esterification by sterol O-acyltransferase is required for lipid droplet crystals to form in Hep3B and RAW264.7 cells and in mice with HFFC diet-induced MASH [33]. Using Raman scattering microscopy, Xu et al. [12] showed that saturated cholesteryl ester spatially correlates with crystals in lipid droplets of livers from humans with MASH. Moreover, in cell models it has been shown that increasing the cholesteryl ester to triglyceride ratio can alter lipid droplet organization and trigger a crystalline state [29–31]. These studies led us to hypothesize that hepatic lipid droplet cholesterol crystals may be composed of saturated cholesteryl ester.

To investigate the hypothesis, we tested if supplementing cholesterol-loaded Hep3B cells with monounsaturated fat acid (MUFA) causes a reduction in lipid droplet cholesterol crystals. Our reasoning was that, under this condition, cells should preferentially produce a higher level of unsaturated cholesteryl esters (e.g., cholesteryl oleate), which in turn was expected to have less propensity to form liquid crystals compared to saturated cholesteryl ester (e.g., cholesteryl palmitate). Indeed, treatment with oleate and palmitoleate (i.e., MUFAs) substantially reduced crystals in Hep3B cells. This effect was not exclusive to Hep3B cells though as oleate also reduced the level of crystals in cholesterol-loaded RAW 264.7 cells. In contrast, treating cholesterol-loaded Hep3B cells with the saturated fatty acid palmitate had no effect on crystals and an inhibitor of the enzyme that converts saturated fat to MUFA increased crystals. Lipidomic analysis profiled the changes in cholesteryl ester species, which would have been contained in lipid droplets. Compared to BSA control, the variable importance in prediction score and univariate analyses showed that Hep3B cells treated with cholesterol had increased saturated (cholesteryl palmitate and cholesteryl stearate) and unsaturated cholesteryl ester (cholesteryl oleate). In contrast, cells treated with cholesterol + oleate had relatively reduced cholesteryl palmitate and cholesteryl stearate and increased cholesteryl oleate compared to the cells treated with cholesterol alone. In other words, the cholesterol + oleate treated cells, which had a small number of crystals, had a higher unsaturated to saturated cholesteryl ester ratio compared to the cholesterol treated cells that had a large number of crystals.

In complement to experiments on cultured cells, we also examined the molecular features of hepatic lipid droplet crystals in liver from mice with diet-induced MASH. In this case, we compared control diet fed mice to mice fed HFFC diet enriched with palmitate and cholesterol, as in previous [13, 33, 39]. HFFC diet feeding increased body and liver weight and caused MASH and the onset of hepatic cholesterol crystals. The biochemical composition of these livers were examined using the advantages of Fourier Transform Infrared (FTIR) spectromicroscopy [42]. Our analyses revealed that, as expected, livers from control diet fed mice had more protein-rich regions whereas livers from HFFC diet fed mice had more lipid-rich regions. Importantly, FTIR spectromicroscopy detected cholesterol crystal-rich regions in the lipid droplets of livers from all of the HFFC diet fed mice but not in any of the livers from Control diet fed mice. Spectra analyses of crystal compared to non-crystal regions as well as reference lipid standards in the spectra corresponding with the ester carbonyl and wagging mode peaks, revealed that hepatic lipid droplet crystals were composed of cholesteryl palmitate, which is consistent with evidence found in livers from humans with MASH [12]. Our investigation support the hypothesis. We identify cholesteryl palmitate as the constituent of crystals in hepatic lipid droplets of HFFC diet fed mice with MASH and our cell culture studies show crystals can be mitigated by increasing the cellular MUFA content, which increased the level of unsaturated cholesteryl ester relative to saturated cholesteryl ester. However, there are other contributing factors that require consideration. One is that MUFA treated cells had greater triglyceride, corresponding with more lipid droplets and several studies demonstrate that an increase in the cholesteryl ester to triglyceride ratio can contribute to the formation of lipid droplet crystals [29–32]. In these studies, however, the levels of saturated versus unsaturated cholesteryl ester and the saturation status of fatty acids within the triglyceride was not examined but may have played a role in the outcomes. Indeed, we found treatments that increase saturated fats had a strikingly different effect on crystals than MUFA supplementation. Studies controlling for these variables may be warranted to disentangle the effects of these distinct neutral lipid species.

It is interesting to consider the effect saturated cholesteryl esters (i.e., cholesteryl palmitate) may have on lipid droplets. The linear structure of cholesteryl palmitate is much more prone to form a crystal at physiological temperatures than a nonlinear cholesteryl oleate, and while cholesteryl oleate forms a type 2 monolayer crystal, cholesteryl palmitate forms a bilayer crystal which may have a distinct effect on lipid droplet physiology [38]. Previous works have shown that a liquid crystal phase transition can alter the lipid droplet proteome and reflect changes in metabolic states that relate to lipid flux [30, 31]. It may even be the case that, in certain conditions, cells regulate lipid composition in lipid droplets to induce crystal formation as a defence response. For example, though they did not assess crystal content, a recent study showed that cholesteryl ester rich lipid droplets associate with inflammatory signaling [50]. New insight regarding these considerations may reveal if and how hepatic lipid droplet cholesterol crystals contribute to MASH as well as more progressive liver pathology. Based on these studies and our results here, we speculate that the high ordering effects of cholesteryl palmitate crystals may reorganize lipid droplets and restrict access to certain metabolic enzymes. Our study also suggests that enhancing the unsaturated cholesteryl ester content may reverse such an effect. Whether or not this contributes to MASH warrants further investigation.

## Abbreviations

ANOVA: analysis of variance
BCA: bicinchoninic acid
DMEM: Dulbecco’s modified eagle medium
DMSO: dimethyl sulfoxide
Hep3B: human hepatoma 3B h hours
MASH: metabolic dysfunction-associated steatohepatitis
MASLD: metabolic dysfunction-associated steatotic liver disease
MßCD: methyl-ß-cyclodextrin
PBS: phosphate buffered saline
SEM: standard error of the mean
SOAT: sterol O-acyltransferase

