## Supplemental Figure 1 for "Monounsaturated fat prevent, whereas saturated fat promote, cholesterol crystals in cytoplasmic lipid droplets"

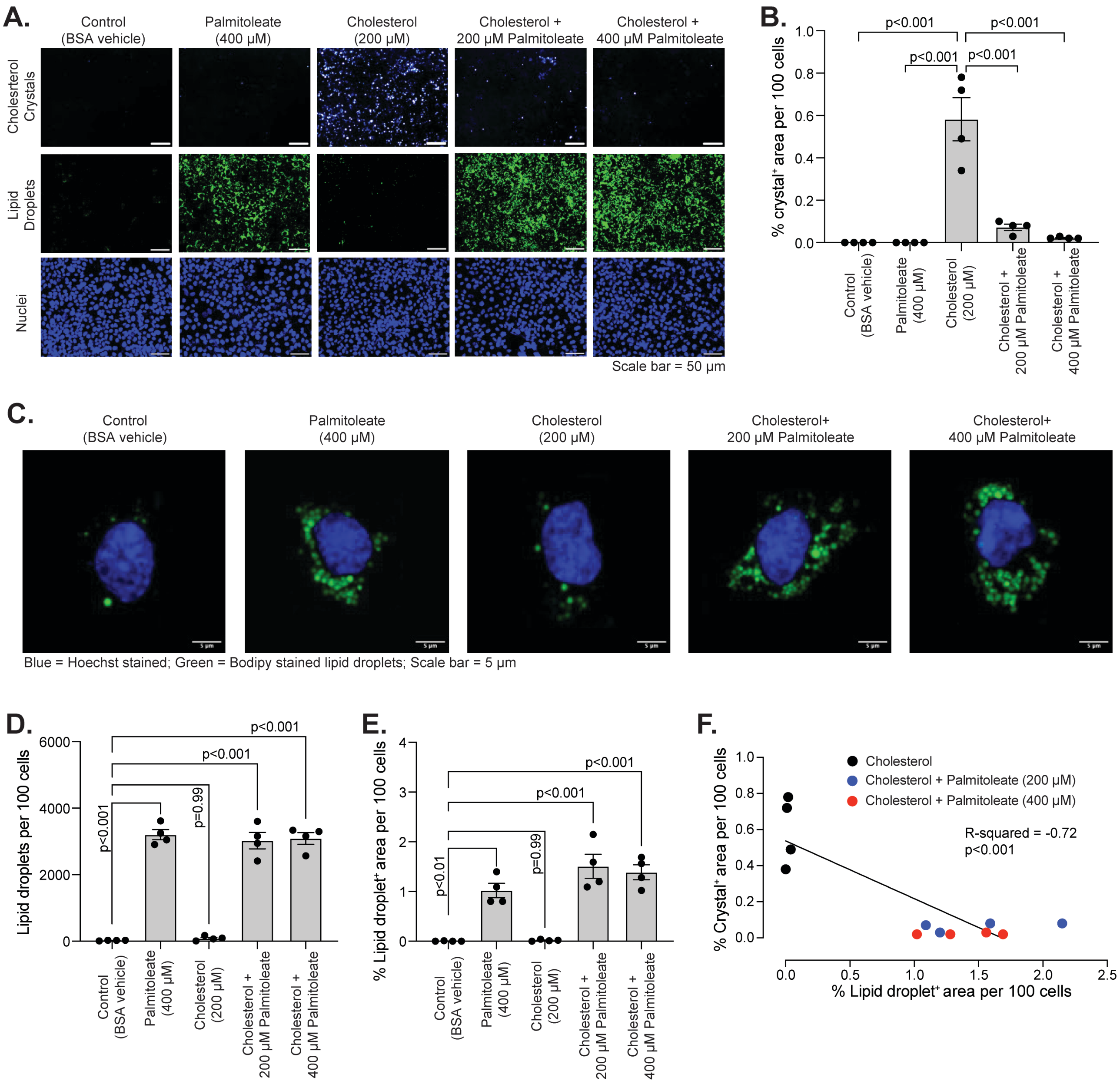


**Supplementary Figure 1**. Supplementation with the monounsaturated fat palmitoleate prevents formation of cholesterol crystals in Hep3B cells. (A) Representative images of cells treated without or with 200 µM cholesterol ± 200 µM or 400 µM palmitoleate. Identified crystals, nuclei, and lipid droplets are shown in panel. (B) Quantified crystals in (A) to examine the effect of palmioleate on cholesterol crystal formation (n = 4). (C) Representative higher magnification images of cells treated as indicated in panel. (D-E) Quantified number (D) and area (E) of lipid droplets (n = 4). Data in B, D, and E are mean ± standard error of the mean and the p-values were determined by one-way analysis of variance, with Tukey's post-hoc test. (F) Correlation analysis of cholesterol crystals with lipid droplets area. The individual data points and slope line are shown (n = 12). The p-value and slope were determined using simple linear regression analysis.
